# Pathogenicity is associated with population structure in a fungal pathogen of humans

**DOI:** 10.1101/2024.07.05.602241

**Authors:** E. Anne Hatmaker, Amelia E. Barber, Milton T. Drott, Thomas J. C. Sauters, Ana Alastruey-Izquierdo, Dea Garcia-Hermoso, Oliver Kurzai, Antonis Rokas

## Abstract

*Aspergillus flavus* is a clinically and agriculturally important saprotrophic fungus responsible for severe human infections and extensive crop losses. We analyzed genomic data from 250 (95 clinical and 155 environmental) *A. flavus* isolates from 9 countries, including 70 newly sequenced clinical isolates, to examine population and pan-genome structure and their relationship to pathogenicity. We identified five *A. flavus* populations, including a new population, D, corresponding to distinct clades in the genome-wide phylogeny. Strikingly, > 75% of clinical isolates were from population D. Accessory genes, including genes within biosynthetic gene clusters, were significantly more common in some populations but rare in others. Population D was enriched for genes associated with zinc ion binding, lipid metabolism, and certain types of hydrolase activity. In contrast to the major human pathogen *Aspergillus fumigatus*, *A. flavus* pathogenicity in humans is strongly associated with population structure, making it a great system for investigating how population-specific genes contribute to pathogenicity.

## Introduction

The fungal genus *Aspergillus* (subphylum Pezizomycotina, phylum Ascomycota) comprises some of the most important human fungal pathogens. Aspergillosis encompasses a range of diseases caused by *Aspergillus* species; these include invasive aspergillosis and chronic pulmonary aspergillosis [1], which impact ∼250,000 and 3 million patients annually on a global scale, respectively [2]. Invasive aspergillosis mainly afflicts individuals with compromised immunity or other underlying conditions [3–7]. Mortality due to invasive aspergillosis varies among patient populations; ICU patients, as well as those with lung cancer, generally exhibit ∼50% mortality [8]. In contrast to invasive aspergillosis, keratitis caused by *Aspergillus* spp. mainly occurs in immunocompetent patients after ocular trauma or contact lens use and can result in visual impairment and even blindness [9]. Fungal keratitis is estimated to cause over one million cases of blindness annually [10].

Molecular barcoding studies from the last decade suggest that the most common infectious agents are *Aspergillus fumigatus* (4-52%), followed by *Aspergillus flavus* (13-40%), *Aspergillus niger* (8-35%), and *Aspergillus terreus* (0-7%) [11–13]. Despite belonging to the same genus, these species exhibit high levels of genomic sequence divergence; for example, *A. fumigatus* and *A. flavus* are as diverged as human and fish genomes [14]. Pathogenicity in *Aspergillus* has evolved independently multiple times, with species following various evolutionary trajectories to become pathogenic [15]. Alongside evolutionary differences, diverse geographic regions exhibit distinct epidemiological patterns in *Aspergillus* species prevalence. For example, the percentage of invasive aspergillosis cases caused by *A. flavus* varies by region, with ∼10% of cases in the USA and Canada [16] but ∼40% in India [13] attributed to *A. flavus*. The incidence of keratitis also exhibits regional differences in etiological agents [9,17].

Despite several *Aspergillus* species infecting humans, research to date has focused primarily on *A. fumigatus*. Unlike *A. fumigatus*, *A. flavus* is of both clinical and agricultural interest [18]. *A. flavus* is notorious for producing the highly carcinogenic mycotoxins known as aflatoxins. Aflatoxins are associated with billion-dollar crop losses annually, and consumption by humans and other animals is associated with cancer, stunted growth, liver failure, and even death [19]. The production of aflatoxins and an arsenal of other small, bioactive molecules termed secondary metabolites varies among populations of the fungus, suggesting niche adaptation to specific microenvironments or competition [20]. In some fungi, including *A. fumigatus*, secondary metabolites like gliotoxin can suppress host immune systems, serving as virulence factors [15,21]. Some strains of *A. flavus* and *A. terreus* also produce gliotoxin or its precursors [22]. While there is some evidence that kojic acid produced by *A. flavus* increases the toxicity of aflatoxin in some insect species [23], to our knowledge, virulence factors that impact human infections have yet to be described for *A. flavus*.

Individual isolates of *A. flavus* exhibit substantial variation or strain heterogeneity. Environmental (i.e., plant- or soil-associated) isolates can cause disease in animal models of aspergillosis [24] and keratitis [25], but strains vary substantially in clinically relevant traits such as growth under iron starvation conditions and virulence in animal models of fungal disease [24,25]. *A. flavus* isolates also vary widely in susceptibility to many antifungal compounds, with the minimum inhibitory concentration of the frontline antifungal, voriconazole, ranging from 0.25-8 µL/mL for isolates from ocular infections [26]. Additionally, *A. flavus* isolates may produce large or small sclerotia (a hardened mass of compacted mycelium capable of long-term survival under stressful conditions), resulting in L and S morphotypes, respectively; sclerotial morphotypes correlate with aflatoxin production and other cellular processes, including conidiation [27].

Although genetic diversity within *A. flavus* has been studied using microsatellite markers in environmental [28–30], veterinary [31], and clinical [32] contexts, these studies focused on a few loci and therefore examine only a small fraction of the total genomic divergence among the isolates. The largest study of *A. flavus* focused on agricultural isolates from the USA, finding high levels of genetic diversity and genetically isolated populations that vary in extent of recombination [33]. However, this study did not include clinical (i.e., human-associated) isolates, and the relationship between environmental and clinical isolates has remained unexplored. Although public databases such as NCBI’s GenBank and Sequence Read Archive contain many *A. flavus* genome assemblies and whole genome sequencing (WGS) datasets, only a small proportion of these are from clinical isolates [26,27]. WGS data provide opportunities to study not only fine-scale population structure (population genomics), but also gene presence and absence across all available genomes in a species (pan-genomics). In a pan-genome, the core genome is defined as those genes found in all or most individuals while those that are missing from some individuals are called accessory genes. The strong conservation of core genes is thought to represent the core metabolism and housekeeping functions of a species [34], with accessory genes encoding non-essential functions and possible local niche adaptations. Pan-genomic analyses are widespread in bacteria but have only recently been adopted in fungi [35]. In *A. fumigatus*, pan-genomes have been used to identify genetic variants associated with human pathogenicity, recombination rates, and similarities between clinical and environmental isolates [36–38].

In this study, we sequenced the genomes of 70 clinical isolates of *A. flavus* representing a diversity of human infection types. We combined the sequencing results with publicly available data to create a dataset of 250 (95 clinical and 155 environmental) genomes. We analyzed the genomes of these isolates to a) infer the population structure of the species, b) investigate the relationship between population structure and isolation environment (clinical vs. environmental), c) define the pan-genome of *A. flavus*, and d) identify genetic elements that are associated with clinical isolates.

## Materials and Methods

### Retrieval of publicly available data

We obtained data for 180 *A. flavus* isolates with paired-end Illumina whole genome sequencing data available on National Center for Biotechnology Information (NCBI) Sequence Read Archive (SRA) in July 2021, including data from 25 clinical isolates. Additionally, *A. flavus* NRRL 3357, which has a chromosome-level genome assembly, was used as a reference. Of the SRA dataset, 152 isolates also had published genome assemblies available through NCBI (Table S1). Eight genomes were from isolates known to produce S-type sclerotia. Isolates represent diverse sources including soil, seed, and plant-associated microenvironments (Table S1).

### Collection of A. flavus clinical isolates and genome sequencing

We also sequenced 70 patient-derived isolates for this study. Of the newly sequenced isolates, 48 were obtained from the German National Reference Center for Invasive Fungal Disease (NRZMyk). Another 15 were from the culture collection of the National Reference Center for Invasive Mycoses and Antifungals (CNRMA) at the Institut Pasteur, France. An additional seven isolates were obtained from the National Centre for Microbiology (CNM) culture collection in Spain. Isolates were from patients diagnosed with keratitis, aspergillosis, and otomycosis and were obtained through a variety of methods (Table S1).

Isolates from the NRZMyk in Germany were grown in Saboraud glucose broth shaken at 37 °C. Species identification was performed using ITS and/or beta-tubulin sequencing. Mycelia were homogenized using a FastPrep (MP Biomedical), and DNA was extracted following the manufacturer’s protocol using the Zymo Research Bacterial/Fungal DNA kit. Isolates from the CNRMA in France were subcultured on 2% malt extract agar (Oxoid) and potato dextrose agar (BD Diagnostic Systems) for 5 days at 30 °C. Species identification was based on macroscopic and microscopic criteria. DNA was extracted as described in Garcia-Hermoso et al. [39]. Briefly, cells were disrupted using the MAGNA Lyser instrument (Roche Diagnostics) with ceramic beads and ATL lysing solution (Qiagen). DNA was then purified using the KingFisher Flex magnetic particle processor system (ThermoFisher Scientific). Isolates from the CNM in Spain were grown in glucose-yeast extract peptone liquid medium with 0.3% yeast extract and 1% peptone (Difco), and 2% glucose (Sigma-Aldrich) for 24-48 hr at 30 °C. Cells were disrupted mechanically by vortex with silica beads. DNA was extracted using the phenol-chloroform method [40].

DNA libraries were prepared using the Nextera DNA Library PrepKit (Illumina, San Diego, CA, USA), according to manufacturer’s guidelines. Sequencing for the isolates from Germany and France (63 total) was performed at Vanderbilt University’s sequencing facility, VANTAGE, using the Illumina NovaSeq 6000 instrument, following manufacturer’s protocols. Sequencing for the seven Spanish CNM isolates was performed using the Illumina MiSeq system, following the manufacturer’s protocols. All sequencing resulted in 150 bp paired-end reads.

### Read mapping and population genomics

By combining the publicly available data from 180 isolates with our newly sequenced clinical isolates, we complied a dataset of 250 *A. flavus* isolates. Draft genome assemblies were available from NCBI for 152 isolates (9 clinical and 143 environmental) [25,33,41–44]. Raw reads from the 70 newly sequenced clinical isolates and 28 publicly available SRA datasets without genome assemblies (16 clinical and 12 environmental isolates) were trimmed using Trimmomatic v0.39 [45] for paired-end data. Trimmed reads were mapped to the NRRL 3357 reference [46] using Bowtie2 v2.3.4.1 with default parameters [47]. We used SAMtools v1.6 [48] to convert the resulting data files to BAM format and sort the BAM files. The AddOrReplaceReadGroups option in Picard tools v2.17.10 (https://broadinstitute.github.io/picard/) was used to append read group labels to BAM files. The Genome Analysis Tool Kit v3.8 (GATK) RealignerTargetCreator and IndelRealigner options were used to produce realigned BAM files [49] and duplicates were removed using the MarkDuplicates option in Picard. We called variants for each genome using the GATK HaplotypeCaller option with -ploidy 1 for haploid organisms. GVCF files were combined using the CombineGVCFs option and the combined file genotyped using the GenotypeGVCFs option. Variants include single nucleotide polymorphisms, insertions, and deletions, so only SNPs were selected and retained. SNPs were filtered using the VariantFiltration option, with *--filter-expression* parameters “QD < 2.0”, “QUAL < 30”, “MQ < 40.0”, “MQRankSum < −12.5”, “SOR >3.0”, “FS > 60.0”, and “ReadPosRankSum < −8.0”; other parameters were set as *--cluster* 8 and *--window* 10, according to the GATK best practices workflow (https://gatk.broadinstitute.org/hc/en-us/articles/360036194592-Getting-started-with-GATK4).

Biallelic SNPs that passed hard filters were retained for further analysis. Biallelic loci refer to loci which at the population level only have two alleles: the reference and an alternative. We identified 9 isolates from the same patient as clones due to their very high genome-wide average nucleotide identity, and 8 of these were excluded from the dataset, leaving 242 isolates. We conducted a principal components analysis in R using adegenet [50]. Adegenet was also used for the discriminant analysis of principal components (DAPC), a multivariate method to determine the optimal number of genetic clusters for a given dataset [51]. The Bayesian Information Criterion (BIC) score was used to evaluate a range of possible numbers of genetic clusters from 1 to 10. The optimal number of clusters was determined by graphing the BIC score for each possible number of clusters. Calculations of missing data per population and sample, minor allele frequency, and Nei’s genetic distance were conducted using the R package SambaR [52]. The R package LEA [53] was used to estimate ancestry coefficients [54], from K = 2 to K = 6. Optimal K (number of populations) was determined using the entropy coefficient method [55].

To determine if molecular variation among populations was larger than variation within populations, we used Nei’s genetic distance [56] to implement an AMOVA (Analysis of Molecular Variance) in R using the R package pegas [57] with 1,000 permutations. A two-way ANOVA implemented in PRISM 10 (Graphpad) was used to establish the relationship between isolate sources (soil, plant-associated, and human-associated) and genetic populations identified by DAPC.

In population genomics, correlations between physical and genetic distance of isolates can impact population structure. As such, we tested for isolation by distance [58] using a Mantel test [59] implemented in the R package dartR [60]. Geographic location for clinical isolates was conservatively estimated by using the coordinates of each isolate’s culture collection. Nei’s genetic distance [56] was used for the genetic distance matrix. For the geographic distance matrix, latitude and longitude was either 1) obtained from previously published data or public metadata from NCBI or 2) estimated from listed hospital location for patient-derived isolates. Isolates without latitude or longitude locations were considered missing data and coded as “NA” in the table.

### Phylogenomics

Using the biallelic SNPs, we reconstructed a phylogeny of the 250 *A. flavus* isolates, with the close relative, *Aspergillus minisclerotigenes* (SRA: SRR12001146), as an outgroup. Only loci present in at least eight isolates were included. We used Lewis ascertainment bias correction to include only variable (non-constant) characters from our SNP data [61]. The phylogeny was built using IQ-Tree v.2.2.2.6 [62] using the ModelFinder Plus [63] (-m MFP) option and 1000 ultra-fast replicates for bootstrapping. The GTR+F+R3 model was chosen by IQ-Tree as the best-fit model according to BIC. The consensus tree was used for visualization. We used iTOL v.6 to visualize and annotate the phylogeny [64]. We also constructed a phylogenetic network using SplitsTreeCE [65] with default parameters to examine the relationships among isolates in a neighbor-net network.

To test whether clinical isolates were randomly distributed across the phylogeny or were more likely to be clustered (that is, whether clinical isolates had a phylogenetic signal), we calculated Fritz and Purvis’s D statistic for binary traits [66] using the R package caper [67].

### Genome assembly and annotation

Genomes were assembled using trimmed reads (described above) for the 70 newly sequenced clinical isolates, as well as 16 additional clinical isolates and 12 environmental isolates from NCBI. Each *de novo* assembly was performed using SPAdes v3.15.0 [68] with default parameters except for k-mer count (set to 21, 33, 55, 77, 99, and 127). For all assemblies, scaffolds were filtered using Funannotate v1.8.10 [69] to remove duplicate sequences and those under 500 bp in length. Scaffolds were masked for repeats using RepeatMasker [70] within Funannotate. Mitochondrial sequences and any bacterial, primate, or viral contaminants identified through routine screening upon submission to NCBI were removed from the genome assemblies. All genomes were evaluated for completeness using BUSCO v4.04 with the Eurotiales database of 4,191 single-copy genes [71].

Gene predictions were generated by Funannotate v1.8.10 using the built-in gene models of *Aspergillus oryzae* (section *Flavi*) as predicted by EVidence Modeler [72], with additional evidence provided in the form of amino acid sequences for proteins from the *A. flavus* NRRL 3357 annotation [46]. To validate the gene-prediction procedure, we compared the new annotation of NRRL 3357 to two recent in-depth annotations [46,73] using OrthoVenn2 [74]. Additional functional annotations were obtained through the “annotate” option within Funannotate that uses InterProScan v5.61.93 [75] with default parameters. Predicted biosynthetic gene clusters (BGCs) were identified using the fungal version of antiSMASH v6.0 [76], with default parameters, and collated into a table format using a custom Python v.3.9 script. For specific clusters of interest, we used BLASTn to confirm the presence or absence of backbone genes as defined by antiSMASH (core biosynthetic genes), e.g., querying the *pksA* nucleotide sequence from the *A. flavus* reference strain NRRL 3357 against all genomes to confirm presence or absence.

### Pan-genome analysis

We identified orthologous proteins using OrthoFinder v2.5.4 [77] in all *A. flavus* genomes that had ≥ 95% completeness of the 4,191 genes in the BUSCO Eurotiales gene set [71], resulting in a dataset of 247 isolates. The core genome was defined as in Lofgren et al. [37] as the set of genes that were present in at least 95% of isolates (in our dataset 236 or more); all other genes were considered part of the accessory genome. A subset of the accessory genome, the “cloud” genome includes orthogroups present in less than 5% of isolates. The presence/absence matrix of the accessory genome was visualized using the R package Complex Heatmap [78]. We created a gene accumulation curve and gene frequency histogram using the R packages vegan [79,80], philentropy [81], and ggplot2 [82]. Using vegan [79], we also calculated a distance matrix for the presence or absence of orthogroups within the accessory genome using Jaccard distance. The distance matrix was then used as input for a principal coordinates analysis (weighted classical multidimensional scaling) to visualize population-level differences in accessory genome content. The accessory genome principal coordinates analysis was visualized using ggplot2 [82]. The alpha for Heap’s law was calculated using the R package micropan [83]. Orthogroups were considered population-specific when absent in all isolates of a particular population but present in > 90% of isolates in other populations, consistent with definitions from Lofgren et al. [37].

Orthogroups were then associated with locus tags and InterPro and gene ontology annotations using a custom Python v3.9 script. Analysis of functional annotation differences among the populations was performed by ANOVA using the number of genes in each isolate’s genome that contained each annotation as input using a custom R script. Statistics and Bonferroni false discovery rate correction were performed using base R v4.3.1. Heatmaps were constructed using the R package Complex Heatmap v2.16.0, and plots were made using ggplot v3.4.4.

We used a phylogenetic generalized least squares (PGLS) analysis as conducted in the R package caper [67] to evaluate whether traits were more likely to be shared by closer relatives in accordance with a Brownian motion model of evolution. PGLS analyses incorporate the phylogenetic relationships between individual data points when examining linear regression of variables [84]. We fit a model of genome size against the number of predicted genes, number of predicted tRNAs, and the number of predicted BGCs using a maximum likelihood estimate of lambda. We also fit a model to explain source (clinical or environmental) using the 10 orthogroups with the most variation in number of genes included in the family.

### Antifungal testing of clinical isolates and cyp51C gene tree

Antifungal susceptibility testing against multiple antifungal compounds was conducted routinely at clinical culture collection sites for all clinical isolates sequenced in this study. Each culture collection used the EUCAST broth microdilution method [85], with slight modification for CNRMA isolates following previously established protocols [39]. Isolates from the German NRZMyk culture collection were tested for susceptibility to itraconazole, voriconazole, posaconazole, isavuconazole, and amphotericin B. Isolates from the CNRMA in France were tested against itraconazole, voriconazole, posaconazole, caspofungin, micafungin, terbinafine, and amphotericin B, as well as isavuconazole for isolates collected more recently. The CNM isolates from Spain were tested for susceptibility to itraconazole, voriconazole, posaconazole, caspofungin, micafungin, anidulafungin, terbinafine, and amphotericin B. We also obtained antifungal susceptibility levels from a small subset of public isolates that had minimum inhibitory concentration (MIC) data available. Susceptibility and resistance to itraconazole and isavuconazole were evaluated based on available guidelines [86,87].

Nucleotide sequences of *cyp51C* were obtained from genome annotations and aligned using MAFFT v.7.407 [88]. A maximum likelihood phylogeny was constructed using IQTree v.2.2.2.6 [62] using 1000 ultra-fast replicates for bootstrapping.

## Results

### Five populations of A. flavus were identified, with clinical isolates overrepresented in one population

For newly sequenced genomes from clinical isolates, short-read sequencing resulted in over 10 million paired-end reads for each of the isolates (Table S2). Trimming resulted in 10,326,680 to 53,093,447 paired reads per isolate (Table S2).

To explore population structure within the species, we analyzed the 909,551 biallelic SNPs that we identified in our dataset. We found evidence of five populations based on admixture (Figure S1) and DAPC analyses (Figure S2). In addition to the previously described A, B, C, and S-type populations, we discovered a new population, D (Figure 1; Table S3). We calculated admixture coefficients for each isolate (Figure 1A), revealing higher levels of admixture in populations A, C, and D than in populations B and S-type. A principal coordinates analysis using Euclidian distance showed considerable overlap between populations A, C, and D, but populations B and S-type were distinct from others, without any overlap (Figure 1B). As with the admixture analysis, the DAPC also provided evidence of five populations in our dataset (Figure 2C), based on BIC scores for each cluster (Figure S1C). The S-type population was the smallest (n = 8) and included only isolates previously confirmed to produce S-type sclerotia. The reference strain, *A. flavus* NRRL 3357, was placed in population A. The designated type strain for the species, *A. flavus* NRRL 1957, was placed in population D.

**Figure 1.**
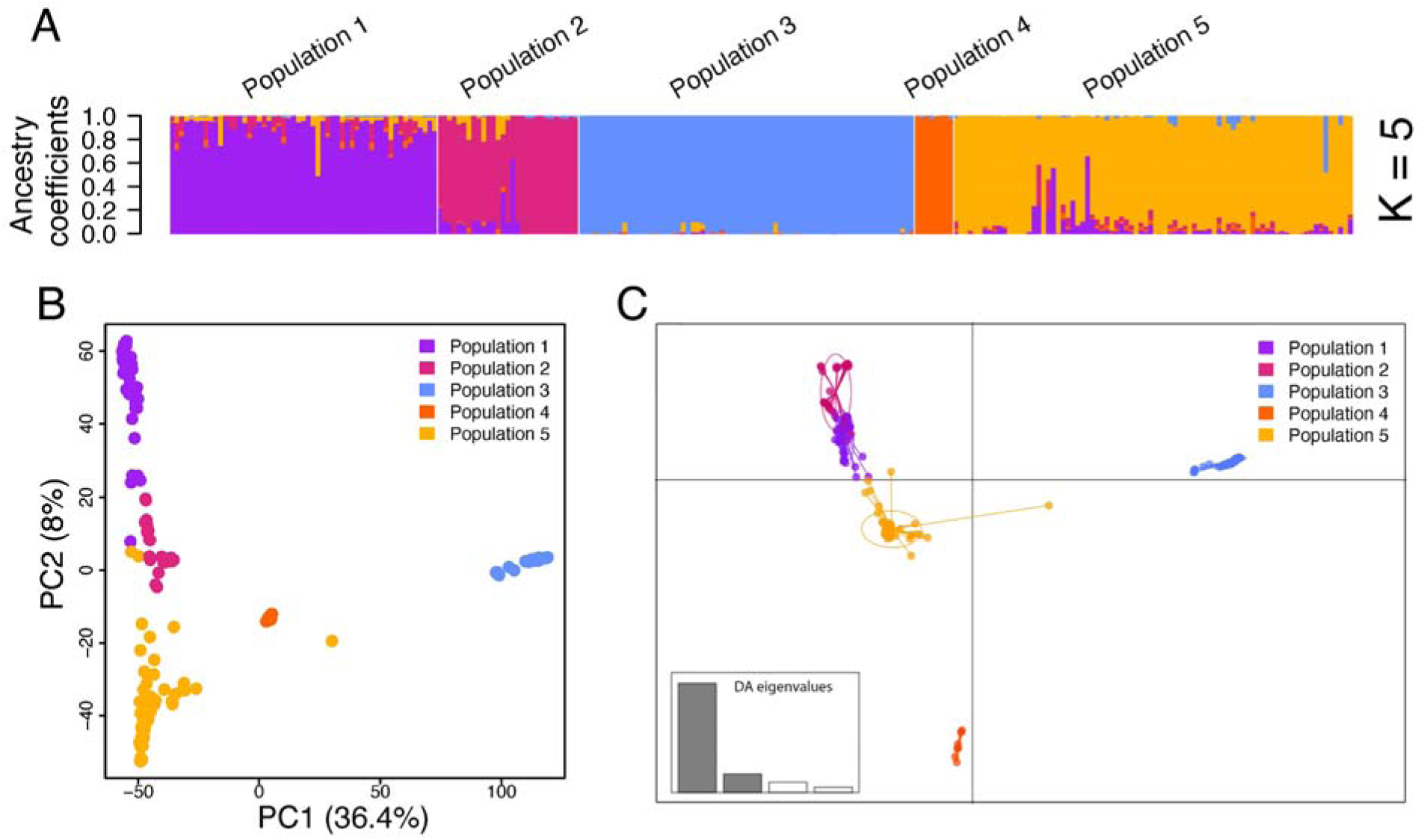
Population structure of *Aspergillus flavus* reveals genetic isolation reflecting five populations, including a new population, D. Analyses are based on 909,551 biallelic single nucleotide variants. A) Estimates of individual ancestry, with K = 5, conducted using the software package LEA [53], which estimates individual admixture coefficients from a genotypic matrix [89]. We estimated admixture for K = 2 through 6, with K = 5 providing the best fit for our data according to the cross-entropy criterion [54,55]. B) A principal coordinates analysis displaying relative genetic distances of individual isolates, here represented by dots, using Nei’s genetic distance matrix. Axes indicate the two principal coordinates retained and the percentage of variance explained by each coordinate. Populations A, C, and D varied primarily along PC2 rather than PC1; population B showed genetic differentiation from all other populations and varied primarily along PC1. C) Discriminant analysis of principal components shows admixture among populations A, C, and D, as well as clear separation of populations B and S-type. Dots represent individuals and ellipses indicate group clustering of individuals. Populations are color coded as indicated in the top right. The discriminant analysis eigenvalues are shown on the bottom left, with the darker bars showing eigenvalues retained.

**Figure 2.**
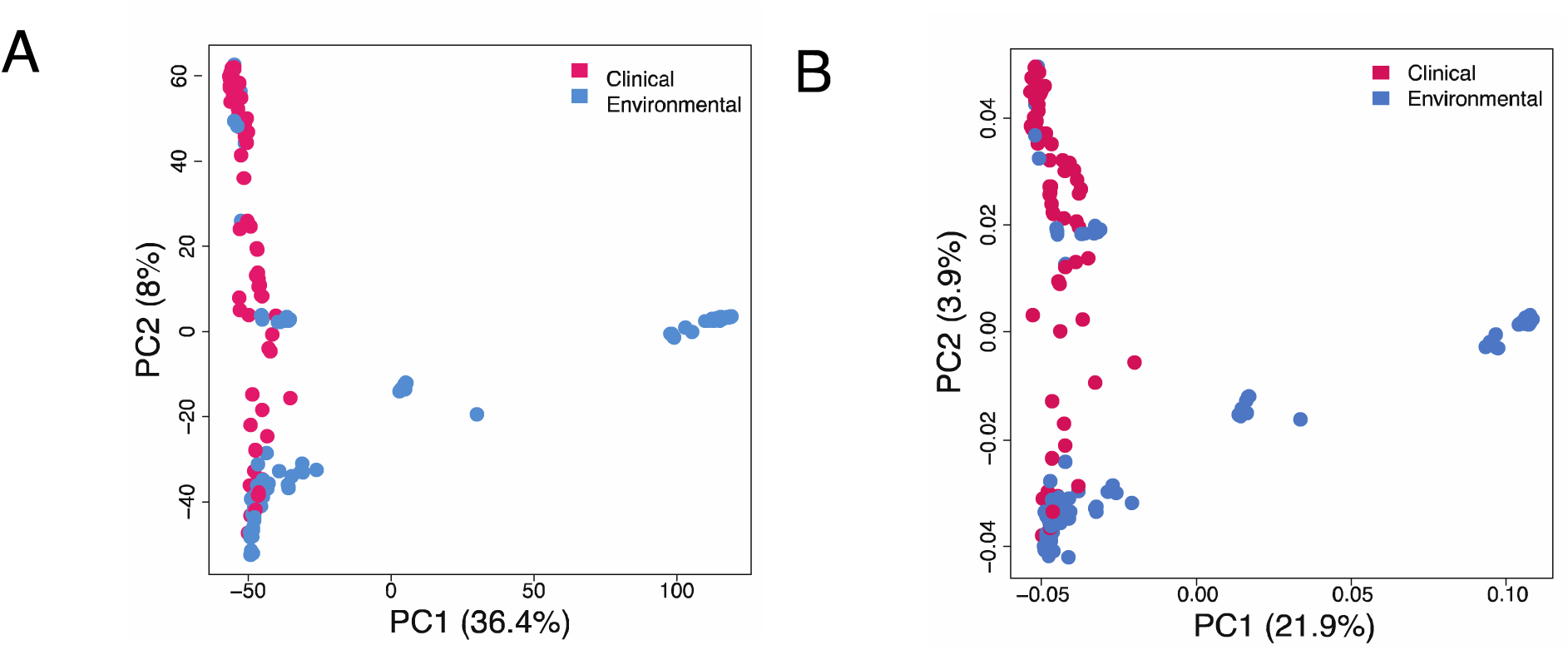
Principal coordinates analysis shows that clinical and environmental isolates of *Aspergillus flavus* are genetically distinct. Each dot represents an individual isolate. Colors indicate the isolation environment of each isolate (clinical or environmental). A) Principal coordinates analysis using Nei’s genetic distance. B) Principal coordinates analysis using Euclidean distance.

The S-type and B populations included almost exclusively isolates from the USA, while the other three populations (A, C, and D) each contained isolates from at least five countries representing three or more continents. To examine the impact of geography on the population structure, we tested for isolation by distance [58]. The Mantel test statistic, or Pearson’s product-moment correlation *r*, lies between −1 and 1, ranging from a perfect negative correlation between the metrics tested and a perfect positive correlation, with 0 indicating no correlation. Although geographic and genetic distance had a marginal positive correlation (Figure S3A), we did not find significant evidence of isolation by distance in the whole dataset (Mantel test; r = 0.4475; p = 0.075). We did, however, see evidence of isolation by distance within populations A, C, and D, but not population B (Figure S3B-E).

Population B also showed the highest level of divergence from other populations, as Nei’s genetic distance was high between population B and all other populations (D was above 0.111 for all comparisons between population B and all others). Nei’s genetic distance was lowest between populations A and D (D = 0.043), indicating genetic similarities between the populations. Genetic differentiation across all five populations was consistent with the results of an AMOVA (Analysis of Molecular Variance) (global phi-statistic = 0.7765; p = 0), indicating more variation among populations than within them.

Populations A, C, and D contained over 95% of the clinical isolates, whereas the S-type population contained exclusively environmental isolates and population B contained only three clinical isolates. Clinical isolates originated from five different countries: India, Japan, Germany, France, and Spain (Table S1). The principal coordinates plots using both Euclidian distance and Nei’s genetic distance differentiated populations with and without clinical isolates along PC1, explaining 36.4% or 20% or the variation, respectively (Figure 2). Both measures of genetic diversity indicate more variation between clinical and environmental isolates than among clinical isolates.

To examine the phylogenetic relationships among the isolates, we next constructed a maximum likelihood phylogeny using the full SNP dataset, with *A. minisclerotigenes* (section *Flavi*) serving as an outgroup (Figure 3). Clinical isolates were in four of the five clades, with each clade corresponding to a genetic population. All isolates with the S-morphotype were placed in a single monophyletic group (Figure 3). Phylogenetic placement of isolates mostly supported the genetic populations (Figure 3). Two clinical isolates were assigned to population A in the DAPC but exhibited considerable admixture with populations A and D (Figure 1A). The maximum likelihood phylogeny recapitulated the DAPC results, with both isolates branching within the population A clade. However, in a neighbor-net network, both isolates were placed within population D rather than population A (Figure S4). Unlike phylogenetic trees, neighbor-net networks capture additional nuance in relationships by including recombination. Based on the admixture analysis and neighbor-net network, these two clinical isolates, one from Germany, and one from India, possibly represent hybridizations between the A and D populations.

**Figure 3.**
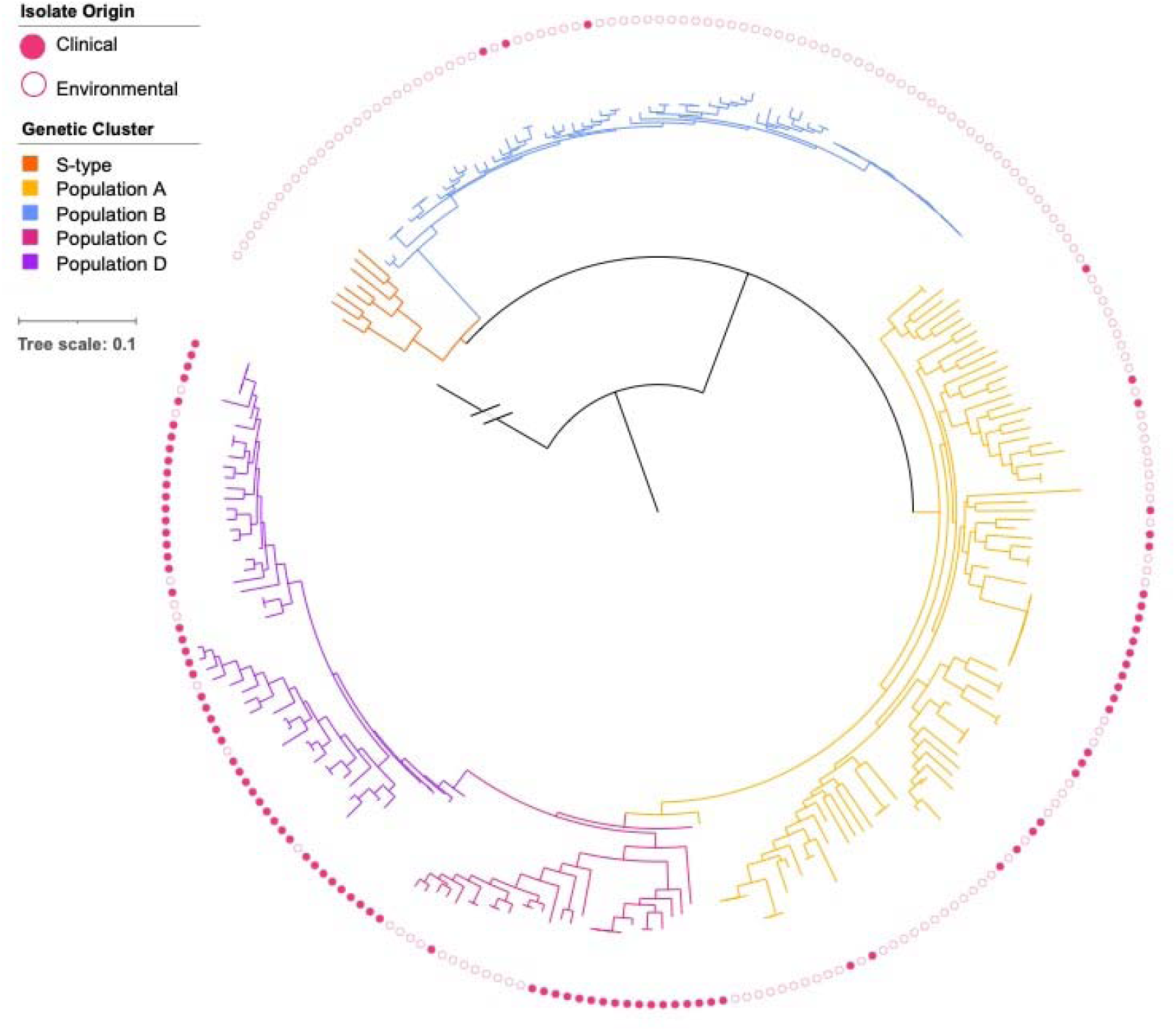
Maximum likelihood phylogeny supports the existence of five populations and non-random distribution of clinical isolates across populations. Filled in circles along the outer track indicate clinical isolates; empty circles indicate environmental isolates. Branch colors correspond with population assignment based on the discriminant analysis of principal components (DAPC; Figure 1): the S-type population is indicated in orange; population A in yellow; population B in blue, population C in pink, and population D in purple. The outgroup, *Aspergillus minisclerotigenes*, is represented in black. Apart from the outgroup, each tip represents an *A. flavus* isolate and branch lengths denote sequence divergence. Out of 131 nodes, 101 had strong support (BS ≥ 95). The phylogeny was constructed using 925,311 SNPs.

To test whether clinical isolates were non-randomly distributed across the *A. flavus* phylogeny, we calculated Fritz and Purvis’s *D* statistic [66]; a value of 0 indicates a clumping of the observed trait (in this case, human pathogenicity) as expected under the Brownian motion model, whereas a value of 1 indicates a random distribution across the phylogeny. The model tests *D* against significant departure from 0 (Brownian motion model of evolution), as well as departure from 1 (random distribution). Consistent with the genetic analyses, we found that *D* was significantly different from a random model indicating clinical isolates were not randomly distributed across the phylogeny (p = 0).

Although we observed enrichment of clinical isolates in some clades, as supported by our *D* statistic, we recognize that our sampling of environmental and clinical isolates was uneven due to hospital culture collection location and availability of public data. All clinical isolates sequenced for this study originate from European culture collections, although we do include public data for additional clinical isolates from India and Japan. Additionally, we do not have any patient-derived isolates from North America, as all our isolates from the USA are environmental.

Despite the uneven sampling, populations with a large proportion of clinical isolates (A, C, and D) include isolates from multiple countries across several continents (Figure S5; Table S1), making it unlikely that the enrichment of clinical isolates stems only from the European provenance of the majority of clinical isolates in the study. If population D, for example, was simply more prevalent in Europe and the overrepresentation in clinical isolates solely due to their abundance in this part of the world, we would expect to see only European isolates within this population, but this is not the case. Roughly half of population D isolates are from Europe; the population includes isolates from seven countries, including clinical isolates from Japan and India, as well as environmental isolates from the USA. In contrast, population B isolates are almost entirely from the USA, although the isolates represent diverse regions within the country. With the available data, we conclude that populations enriched in clinical isolates are more globally widespread than populations lacking clinical isolates (S-type and B populations), although this finding warrants further testing through sequencing of isolates from additional geographic regions.

### A. flavus isolates exhibit heterogeneity in gene content and genome size

Strain heterogeneity in gene content can impact diverse traits, including virulence and secondary metabolite production, so we also examined our dataset in using methods independent of the reference genome. Genomes from the newly sequenced clinical isolates contained 17 to 4,235 scaffolds (Table S2). We also assembled genomes for an additional 16 clinical and 12 environmental isolates from publicly available sequencing data, resulting in genomes containing 700 to 3,615 scaffolds (Table S2). Publicly available genome assemblies of 152 environmental *A. flavus* isolates contained 8 to 1,821 scaffolds [33,42–44,90–93] (Table S1). BUSCO analysis of the genomes confirmed the high completeness (> 95%) of the assemblies for all but three isolates, which were excluded from the pan-genome analysis (Table S4).

We examined variability in genome size among populations, which could indicate genetic expansions or streamlining. The mean genome size of population D was higher than both populations A and B (Tukey’s multiple comparisons test, adjusted p = 0.0175), with all other pairwise comparisons of population means being nonsignificant (Figure S6).

We annotated all genomes using Funannotate [69] to obtain consistent annotations for comparison. To ensure our annotation pipeline resulted in high-quality proteomes, we compared the Funannotate predicted proteome of NRRL 3357 to the RefSeq reference annotation and an additional transcriptome-based annotation of the same strain [73]; the two published proteomes contained 11 and 66 orthogroups that were not present in the Funannotate prediction, respectively, accounting for a tiny fraction of the gene content. Minor differences in gene prediction are expected due to gene fragmentation contributing to orthogroup variation. Overall, we are confident that the annotations predicted by the Funannotate pipeline are consistently high quality, enabling comparisons across isolates. The number of protein-coding genes predicted by Funannotate ranged from 11,461 to 15,501 across isolates (Table S5).

### The A. flavus pan-genome is closed and contains 17,676 orthogroups

To quantify the degree of gene presence-absence variation among isolates, we constructed a pan-genome of *A. flavus.* We used OrthoFinder to cluster predicted proteins into orthogroups, which were then compared across isolates and populations. Our pan-genome of *A. flavus* is closed (Heap’s law, alpha = 1.000023), with each genome after the 200th adding fewer orthogroups (Figure 4A). We identified a total of 17,676 orthogroups. Of these, 10,161 (57.5%) orthogroups were in at least 95% of isolates; we consider these orthogroups to be the core genome. Within the core genome, 3,375 orthogroups were single-copy and present in all isolates. The pan-genome of *A. flavus* exhibits a U-shaped distribution, as expected (Figure 4B). The accessory pan-genome of *A. flavus* consists of 7,515 orthogroups (Figure 4C), of which 3,387 (19.1% of all orthogroups) were in fewer than 5% of isolates and which we consider the “cloud” genome.

**Figure 4.**
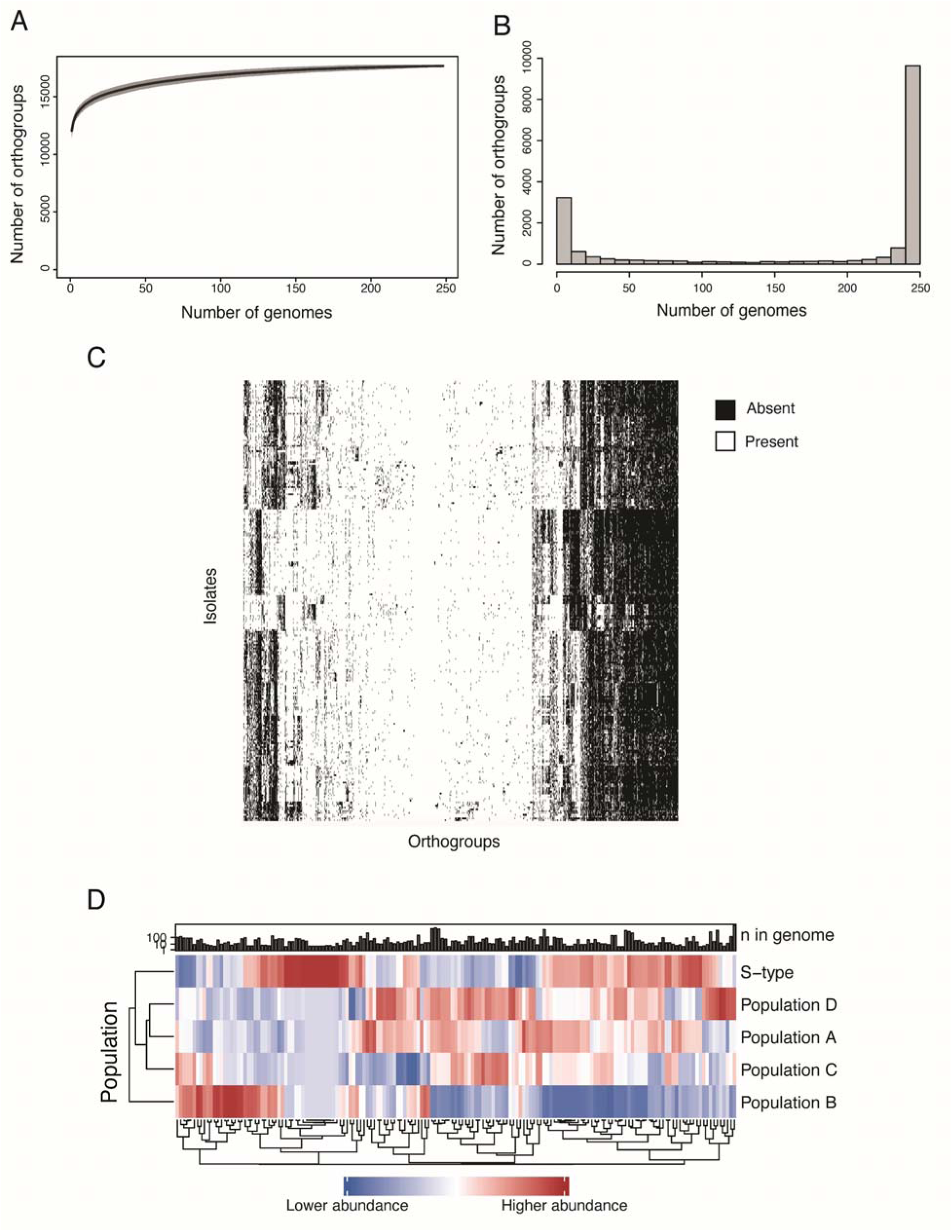
The pan-genome of *Aspergillus flavus* is closed and contains 17,676 orthogroups. A) Rarefaction curve of number of orthogroups added with each additional genome, excluding singletons. B) Histogram of orthogroup frequency determined by number of genomes in which each orthogroups is present. The core genome contains 10,161 orthogroups. The accessory genome of *A. flavus* contains 7,515 orthogroups. C) Presence/absence heatmap of accessory orthogroups. D) Heatmap of abundance of gene ontology (GO) annotations with significant differences in abundance among populations. Significance determined by one-way ANOVA. Bonferroni-corrected, p < 0.05.

To explore which functional annotations were over or underrepresented within populations, we examined presence or absence and abundance of InterPro annotations and gene ontology (GO) terms. Populations A, C, and D, which are enriched in clinical isolates, shared much of their gene content and did not show any population-specific patterning of orthogroup presence or absence in a PCoA, but population B, did (Figure S7). We infer that gene content among population B isolates is more conserved and distinctive from other populations, likely due to low diversity within the population. In addition, we examined the abundance of GO terms and InterPro annotations and compared the mean among populations. Populations had substantial differences in annotations and several GO terms were differentially abundant among populations (Figure 4D). Given the over-representation of clinical isolates in population D, we focused on interpreting differences in functional annotations between population D and all other populations. Isolates in this population had a higher abundance of genes involved in many cellular processes, including certain types of hydrolase activity, nucleoside metabolic and carbohydrate metabolic processes, DNA-binding transcription factor activity, zinc ion binding, regulation of transcription, lipid metabolic process, NAD binding, catalytic activity, and acyltransferase activity (Table S6; Figures 5 and S8). Genes annotated with ferric iron binding functionalities were found in lower abundance compared to other populations.

**Figure 5.**
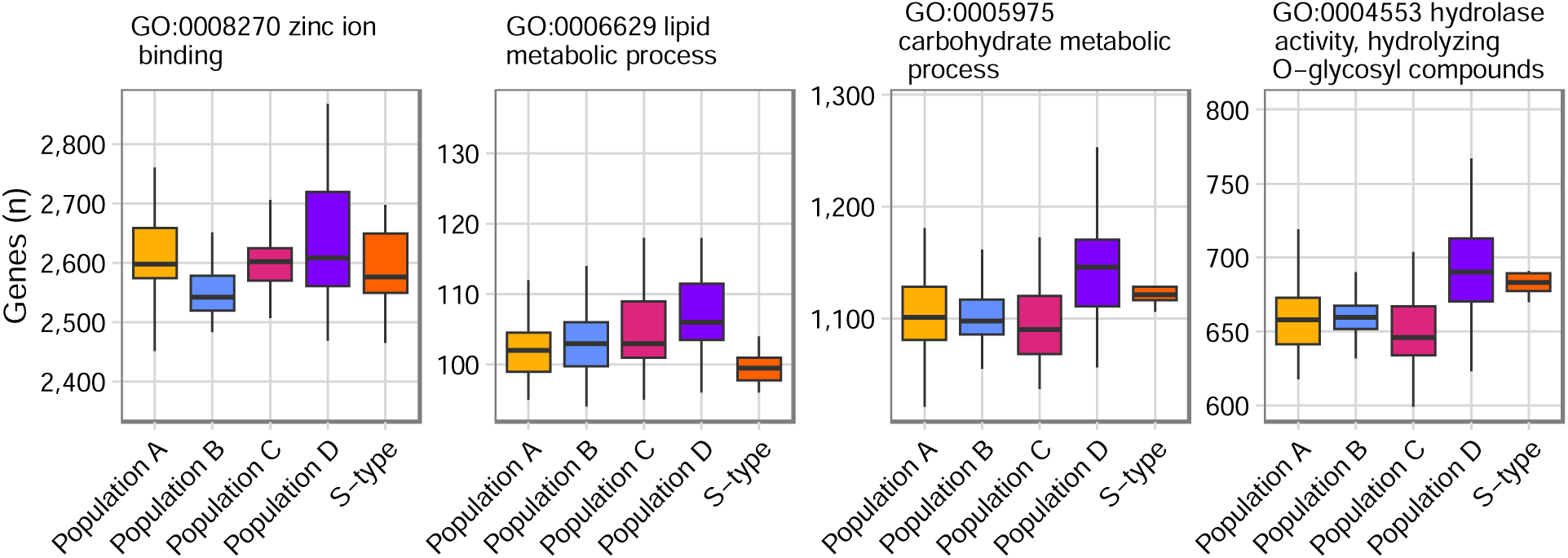
Gene ontology (GO) terms more prevalent in population D than other populations include zinc ion binding and hydrolase activity, among others. Boxplots indicate the number of genes annotated with each GO term per isolate, by population assignment. The Y axis scale is adjusted for each GO term to better show differences among populations.

Genes in a putative non-ribosomal peptide synthesis (NRPS) BGC with an unknown product were absent in > 90% of isolates within the S-type and B populations. The backbone gene for the NRPS cluster was present in all isolates in all populations, but additional biosynthetic or transporter genes within the BGC were absent from isolates within the S-type and B populations (G4B84_009247, G4B84_009246, G4B84_009245, and G4B84_009244 in *A. flavus* NRRL 3357, where all genes in the BGC are present). The BGC was previously identified as “BGC_44” on chromosome 6 but was not explored in depth [20]. GO terms associated with multiple genes within the BGC (OG0011498 [GO:0000981; GO:0006355; GO:0008270]; OG0011868 [GO:0003824, GO:0006807]; OG0011918 [GO:0003824]) were also differentially abundant among populations (Table S6).

Additionally, orthogroups related to aflatoxin biosynthesis (GO:0045122) were abundant at different levels among populations (Table S6). Predictions from antiSMASH indicate the aflatoxin BGC was present in 66.4% of isolates (n = 166). In line with previous research [20], the BGC was present in almost all isolates in populations A, C, and S-type, but absent or degraded in many isolates within population B (Figure S9). Interestingly, we found that the aflatoxin BGC was also absent or degraded in the newly defined population D, which has the most clinical isolates of any population.

### Three orthogroups with high variability in number of gene family members were correlated with human-association

We used phylogenetic generalized least squares (PGLS) models to examine the relationships between multiple variables including isolate source (clinical or environmental), genome size, number of tRNAs, number of predicted genes, and number of predicted biosynthetic clusters. Using these phylogenetically informed linear regression models, we observed a correlation between genome size and number of predicted genes (p = 0; adjusted R^2^ = 0.4891). We also examined orthogroups with the largest variability of gene family members from each isolate. Of the 10 orthogroups examined, only 3 were significantly associated with isolate source (OG0000011, OG0000060, and OG0000270), all with low adjusted R-squared values (Table S6); BLASTp results linked OG0000060 and OG0000270 with hypothetical proteins and implicated OG0000011 in natural product biosynthesis. OG0000011 was annotated with gene ontology terms “GO:0003824 catalytic activity” and “GO:0016746 transferase activity, transferring acyl groups,” which were both significantly differentially abundant among populations (Table S7). None of the PGLS models showed significant correlation between orthogroups and DAPC population assignment (Table S3).

### In vitro antifungal susceptibility testing reveals low levels of azole resistance

Minimum inhibitory concentration (MIC) data for at least one antifungal drug were available for 83 isolates used in this study (Table S8; Figure S8). Breakpoints for resistance have been defined for *A. flavus* for itraconazole and isavuconazole [86]; among the newly sequenced clinical isolates tested, three were resistant to itraconazole (MIC > 1), and two were resistant to isavuconazole (MIC > 2). Additionally, four isolates had MICs above the epidemiological cut-off values (ECOFF) for voriconazole (MIC > 2) and five for posaconazole (MIC > 0.5). The MIC of newly sequenced clinical isolates ranged from 0.06 to 4 for itraconazole, 0.125 to 8 for voriconazole, and ≤ 0.016 to 0.5 for posaconazole (Table S7), with 58 isolates (84%) having MICs below the ECOFF for all three azoles. The MIC for amphotericin B ranged from 0.5 to >16, with 15 of the isolates above ECOFF values, a rate of 21.7% (Table S7); however, *A. flavus* is not considered a good target for amphotericin B and resistance remains undefined [86]. The phylogeny of *cyp51* sequences did not indicate any association between variability in the gene and susceptibility to voriconazole.

Our ability to test population-level differences in fungicide susceptibility was precluded by missing MIC data for almost all environmental isolates and some clinical isolates, as well as the inaccessibility of fungal culture for isolates whose data were retrieved from public databases. Overall, resistance to amphotericin B and voriconazole were low and susceptibility was distributed across the phylogeny (Table S8, Figure S10).

## Discussion

In this study, we examined the population structure and pan-genome of *A. flavus* using genomic data from 250 isolates, including 70 newly sequenced clinical isolates. To our knowledge, this study is the first to combine genomic data for both clinical and environmental isolates of this pathogen, providing a rich dataset for future study and revealing fine-scale differences in pathogenicity within *A. flavus*.

Previous research using genomic data of environmental isolates from the USA described three populations of L-type (isolates producing large sclerotia) *A. flavus* isolates: A, B, and C [33]. With the inclusion of additional isolates, notably clinical isolates, our study identifies another distinct population, here termed population D, which contains the majority of clinical isolates. Clinical isolates were present in all L-type populations, but at different levels— populations A, C, and D were enriched in clinical isolates, with population D containing the highest proportion. Isolates with the small sclerotial morphotype (S-type) grouped together in all analyses, as seen previously [33], and did not include any clinical isolates. We did not expect any clinical isolates to be part of the S-type population, as S-type isolates produce conidia at a far lower rate than L-type isolates [27]—interaction with these airborne asexual spores is how most patients are infected with *Aspergillus* species, so isolates producing fewer conidia are less likely to have spores interact with a human host.

Populations contained a combination of isolates from several different infections or microenvironments including soil, infections of different plant hosts (e.g., peanuts, corn, almonds), and different types of human infections (keratitis, aspergillosis, otomycosis, etc.), indicating a lack of specialization in populations. Previous work on environmental isolates indicated a lack of host specialization in *A. flavus*, with a single isolate able to infect both plant and animal hosts [94], which is consistent with our observation of the lack of clustering of isolates from the same microenvironment.

Interestingly, clinical isolates were concentrated in populations A, C, and D, with population D containing the majority of clinical isolates and few environmental ones. *Cryptococcus neoformans*, another opportunistic fungal pathogen of humans, shows similar enrichments of clinical isolates in some clades compared to others [95], whereas the most common human pathogen in the genus *Aspergillus*, *A. fumigatus*, does not [36]. In *A. flavus* populations A, C, and D, isolates did not cluster by country of origin, but we did observe a marginal positive correlation between genetic and geographic distances, indicating that genetically divergent isolates were likely to be geographically distant. Geographic sampling within our dataset was not balanced, due to our use of public data and our lack of access to clinical isolates outside of Europe.

We observed several important differences between *A. flavus* and the major human pathogen *A. fumigatus*. Our finding that some populations are highly enriched for clinical isolates in *A. flavus* contrasts from observations in *A. fumigatus*, wherein clinical isolates are more evenly distributed across all clades [36], highlighting the importance of studying *A. flavus* as a pathogen rather than assuming that pathogenicity in the two species evolved similarly. Ecological differences among species contribute to the various clinical presentations and prevalence of *Aspergillus* species causing aspergillosis [96], such as the ability to form biofilms or the size of conidia. Likewise, genetic differences among species may explain some of the variance and prevalence of *A. flavus* related to *A. fumigatus*. *A. flavus* has a larger pan-genome than *A. fumigatus*, likely due to the difference in genome size between the species, with accessory genes composing a similar percentage of the pan-genome. Several pan-genomic studies have focused on *A. fumigatus*, with the core genome ranging from 55% to 69% of the pan-genome [36–38], compared to our finding of 57% in *A. flavus*. A recent review stated that *A. flavus* clinical isolates were more similar to one another and exhibited lower diversity than clinical isolates of *A. fumigatus*, which were more genetically diverse [97]. *A. fumigatus* has many more genomes available from clinical isolates than *A. flavus* and the isolates are not associated with population structure as in *A. flavus.* However, our observation that clinical isolates are constrained to populations A, C, and D, which share genetic similarities and overlap in a principal coordinates analysis, supports a level of similarity among *A. flavus* clinical isolates despite deep phylogenetic divergences. We advocate for additional sequencing of clinical isolates, particularly from North and South America and Africa, as no genomes of clinical isolates are currently available from these regions. Nevertheless, our analyses suggest that the core genome of *A. flavus* will remain similar even with the addition of new data; among the pan-genomic studies of *A. fumigatus* the core genome was consistent whereas the accessory genome varied based on input data [36–38].

Within the pan-genome of *A. flavus*, we observed several differences among populations. Several GO terms were enriched in population D, often in biological processes like carbohydrate, nucleoside, and lipid metabolism. Molecular functions enriched in population D included hydrolyzing O-glycosyl compounds, a function of exo-polygalacturonases, which are involved in the degradation of plant cell wall polysaccharides [98]. None of the GO terms were directly implicated in functions typically related to pathogenicity, but several GO terms associated with metal acquisition were differentially abundant among populations. For example, zinc ion binding was enriched in population D. Zinc is an essential micronutrient for many fungal processes, and in *A. fumigatus*, deletion of a zinc acquisition factor attenuated virulence in a mouse model of aspergillosis [99]; however, we do not yet know whether the abundance of genes annotated to involve zinc ion binding would confer higher virulence in population D of *A. flavus*.

In other fungal infections of humans, secondary metabolites have been implicated in virulence [100], most notably the role of gliotoxin in *A. fumigatus* infections. However, no secondary metabolites have been associated with *A. flavus* human infections. The most famous secondary metabolite produced by *A. flavus* is aflatoxin. The secondary metabolite is not thought to be important for human infections as the optimum temperature for aflatoxin production is below 37 °C, with transcription of the BGC dropping at higher temperatures [101]. The predicted BGC for aflatoxin follows previously reported population-specific patterns [20], with presence or absence of the aflatoxin genes explained by clade and population.

Although both clinical and environmental isolates within populations A and C contained the aflatoxin BGC, isolates in population D often lacked genes related to aflatoxin biosynthesis. Population B, which included almost entirely environmental isolates, also lacked aflatoxin biosynthesis genes. Hospitals do not measure aflatoxin production for clinical isolates, leading to a paucity of production data for clinical isolates. However, it appears that although some clinical isolates in populations A and C may have the potential to produce aflatoxin, many clinical isolates in population D lack the necessary genes, and we expect them to be non-aflatoxigenic. The absence of the aflatoxin BGC within many clinical isolates of *A. flavus*, including almost all within population D, reinforces the apparent lack of association between aflatoxin and virulence in the context of human infections. Other predicted biosynthetic genes and gene clusters, such as BGC_44 [20], which had accessory biosynthetic genes more prevalent in population D than in other populations, have not been connected to metabolites and therefore their potential role in infection remains unknown.

Resistance to antifungal drugs in human pathogenic fungi continues to be a growing concern [102]. Our examination of susceptibility to multiple antifungal compounds revealed similar ranges of minimum inhibitory concentration (MIC) as seen previously but, compared to prior studies [103–105], our newly sequenced European clinical isolates had a lower range of MICs for itraconazole and higher range for voriconazole. We observed a slightly lower rate of isolates non-susceptible to amphotericin B (21.7%) than seen in environmental isolates in Vietnam (25.7%), and we observed a much lower rate of resistance to azoles [105]. Agricultural fungicide usage has been implicated in *A. fumigatus* resistance to azoles in Europe and linked to specific genetic variants [106], but no similar study has been conducted for *A. flavus*. However, some areas of southeast Asia have high rates of azole resistance in environmental isolates, although resistance was not directly linked to agricultural azole use [105]. As our newly sequenced clinical isolates all originated in Europe, variation in azole susceptibility rates is possibly due to the differences in regulation and usage of triazole fungicides between regions. Mutations within the gene *cyp51C* have been implicated in voriconazole resistance in *A. flavus* [107]. In our dataset however, mutations relative to “wildtype” *cyp51C* did not correlate with higher MIC values, as many susceptible strains had identical nucleotide sequences as strains with known resistance. Other genetic mechanisms of resistance to voriconazole have recently been elucidated, including transient copy-number variation and large-scale segmental duplications of multiple chromosomes [108], which we would not capture in our study.

In summary, we present evidence that clinical isolates of *A. flavus* share genetic similarities and are concentrated in certain populations rather than distributed across the phylogeny, particularly in an apparently non-aflatoxigenic, newly defined clade we have named population D. Clinical isolates from many countries and infection types present in population D. We acknowledge that sampling was uneven and did not cover the full distribution of *A. flavus*, and advocate for additional sampling from regions underrepresented in this dataset. Additionally, accessory genes and the aflatoxin BGC differ between populations, possibly providing future opportunities for distinct agricultural and clinical treatments. Although we did not discover a single genetic element which could explain the difference between clinical and environmental isolates of *A. flavus*, we did discover a new clade of *A. flavus*, enriched in clinical isolates, with distinct genetic features. This *A. flavus* genomic dataset and pan-genome provide a valuable tool for understanding the molecular mechanisms by which some, but not all, isolates of *A. flavus* can cause serious human infections.

## Data availability

Data associated with the newly sequenced genomes from clinical isolates as part of this study, including paired-end reads and draft genome assemblies, are available under BioProject PRJNA836245.

