## Supplementary Figures for "Pathogenicity is associated with population structure in a fungal pathogen of humans"

Supplemental figures for “Pathogenicity is associated with population structure in a fungal pathogen of humans” Hatmaker et al.

S1. Output for admixture analysis performed using LEA.

S2. Optimization of discriminants analysis of principal components.

S3. Isolation by distance using the Mantel test reveals positive correlation.

S4. Neighbor net network of *A. flavus* isolates based on SNPs.

S5. Maximum likelihood phylogeny of *A. flavus* with isolate country of origin indicated.

S6. Genome size of *A. flavus* isolates by population.

S7. Principal coordinates analysis of orthogroups from the accessory genome of *A. flavus*.

S8. Boxplots of GO terms more prevalent in population D than other populations.

S9. Aflatoxin production and biosynthetic gene cluster presence across phylogeny of *A. flavus*.

S10. Cladogram with antifungal susceptibility for voriconazole and amphotericin B.

Figure S1. Output for admixture analysis performed using LEA. A) Admixture plots for each value of K (number of clusters), from 2 to 6. B) Cross-entropy criterion score for each value of K.

**A**

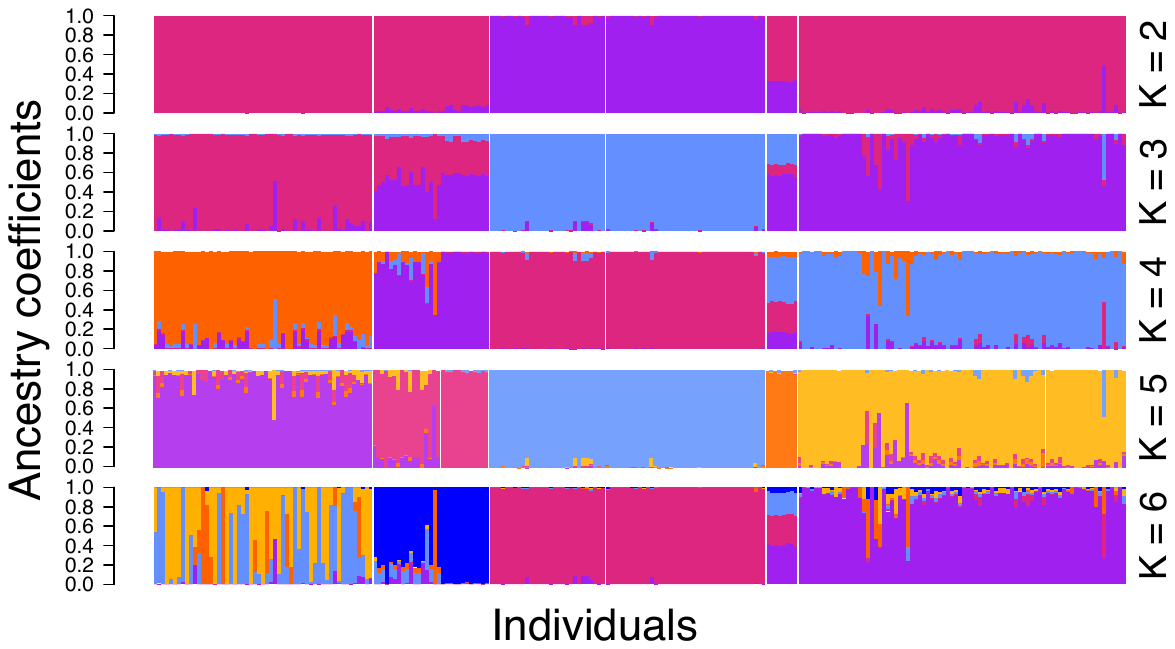

**B**

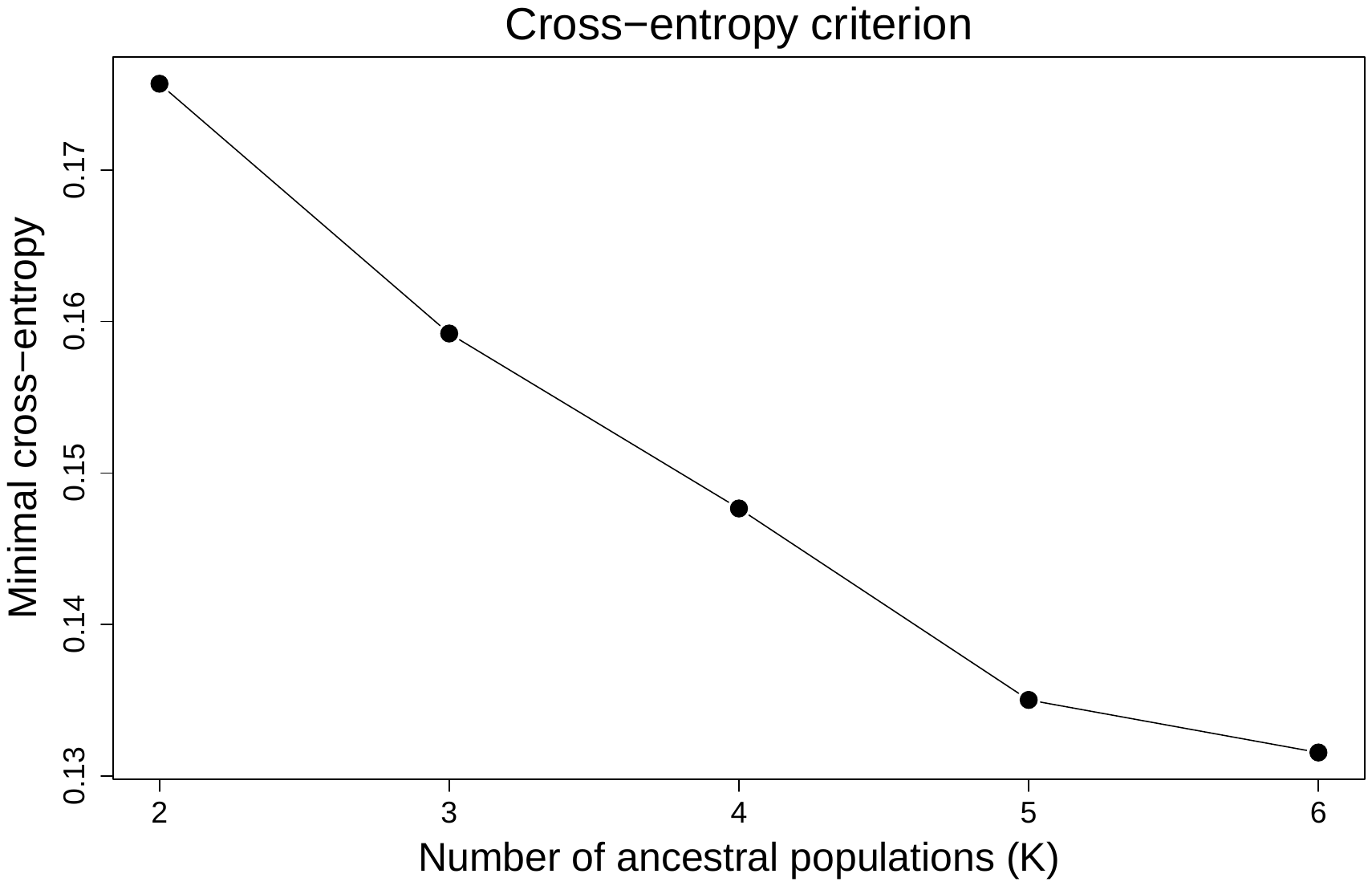

Figure S2. Optimization of discriminants analysis of principal components (DAPC). A) Determination of the optimal number of principal components to retain for the DAPC. B) Contributions of eigenvalues for the DAPC. C) Bayesian Information Criterion score for each value of K (number of clusters).

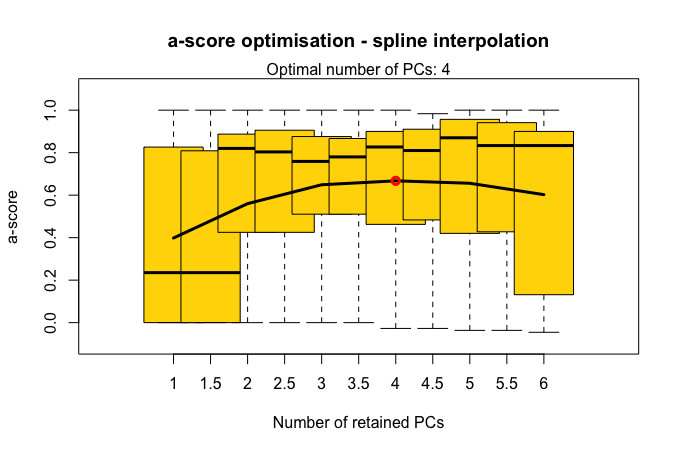
**A**

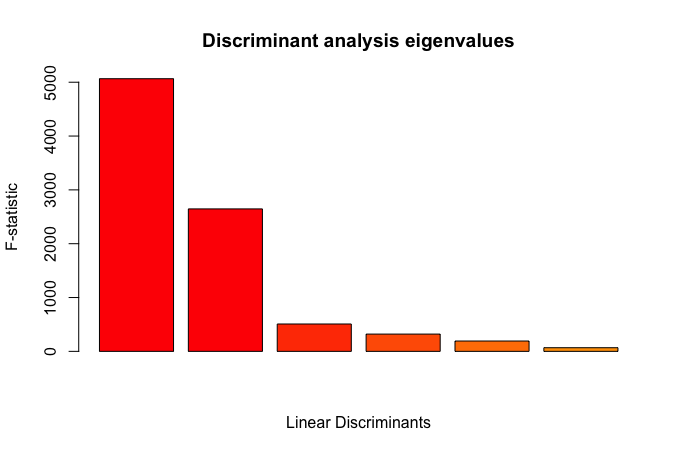
**B**

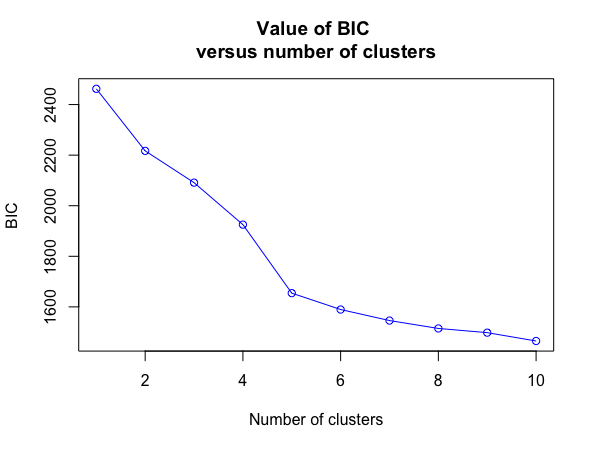
**C**

Figure S3. Isolation by distance using the Mantel test reveals positive correlation for

A) all populations (r = 0.4475; p = 0.075); B) Population A (r = 0.1584; p = 0.001); C) Population C (r = 0.3183; p = 0.001); D) Population D (r = 0.1718; p = 0.018); and E) Population B (not significant, p = 0.066).

**A**

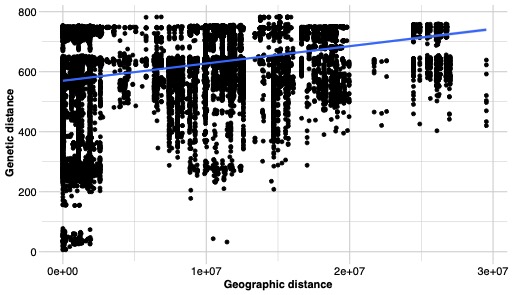

**B	C**

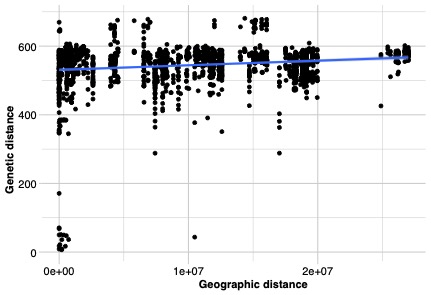

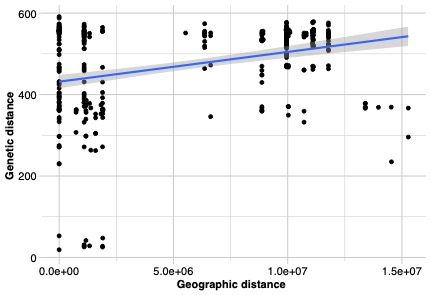

**D	E**

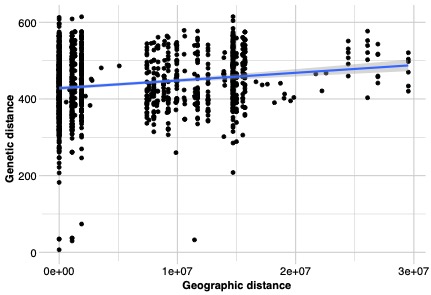

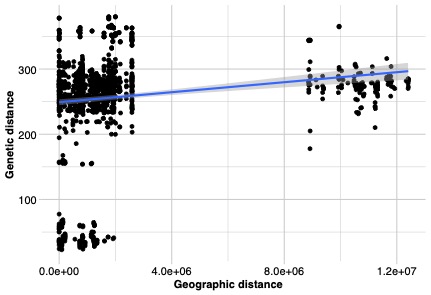

Figure S4. Neighbor net network of *A. flavus* isolates based on SNPs, analyzed and visualized in SplitsTreeCE. Population assignments from the discriminant analysis of principal components are indicated by the label and color of the ellipses. Green stars indicate isolates exhibiting high admixture and assigned to a different population than seen in the neighbor net network.

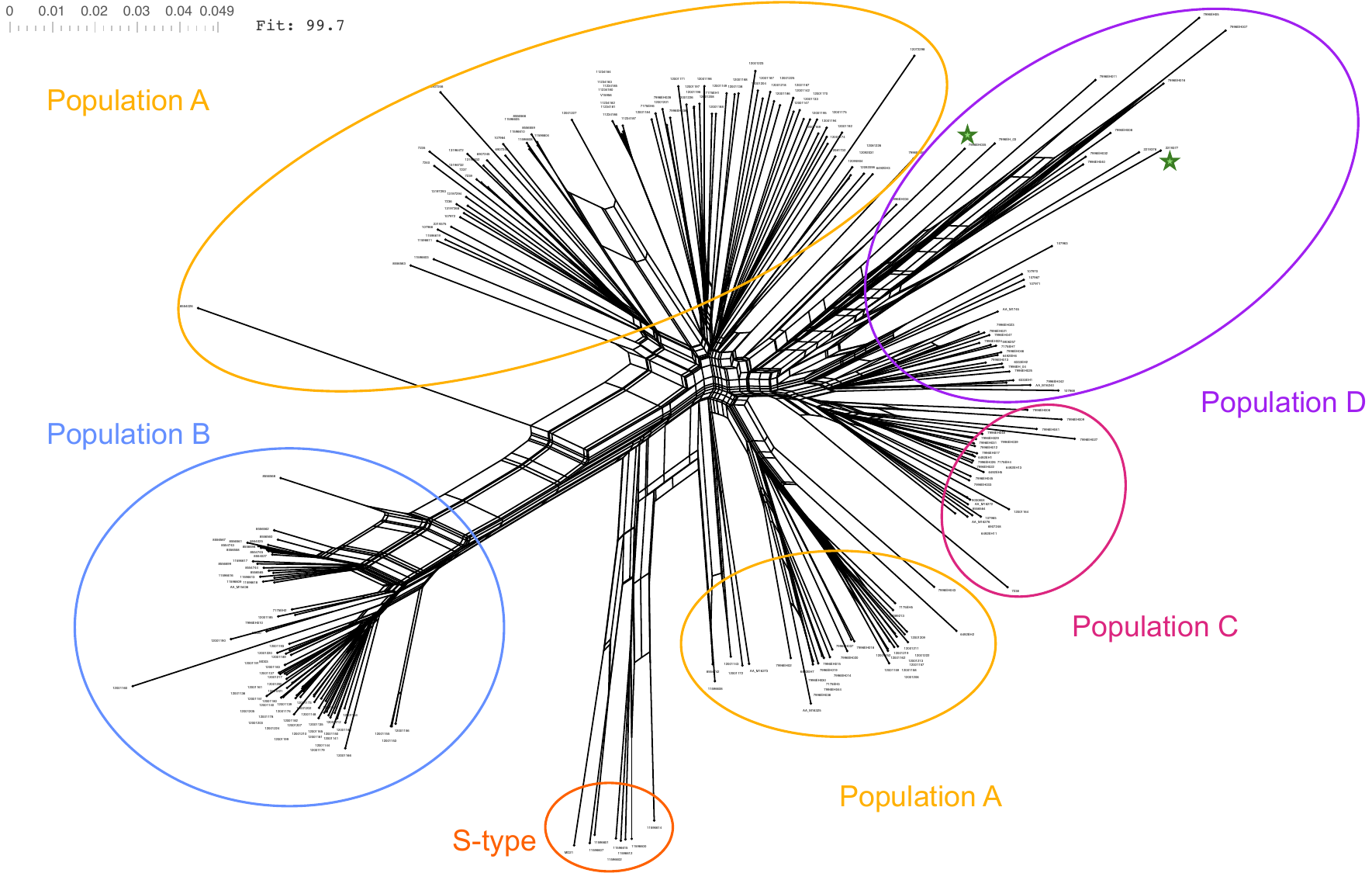

Figure S5. Maximum likelihood phylogeny of *A. flavus* with isolate country of origin indicated on the outer track. Clinical isolates are indicated with filled in pink circles on the inner track. Branches are color-coded based on population assignment from discriminant analysis of principal components. Countries: ETH = Ethiopia, FRA = France, GER = Germany, IND = India, JAP = Japan, NED = Netherlands, PAK = Pakistan, SPA = Spain, USA = United States of America.

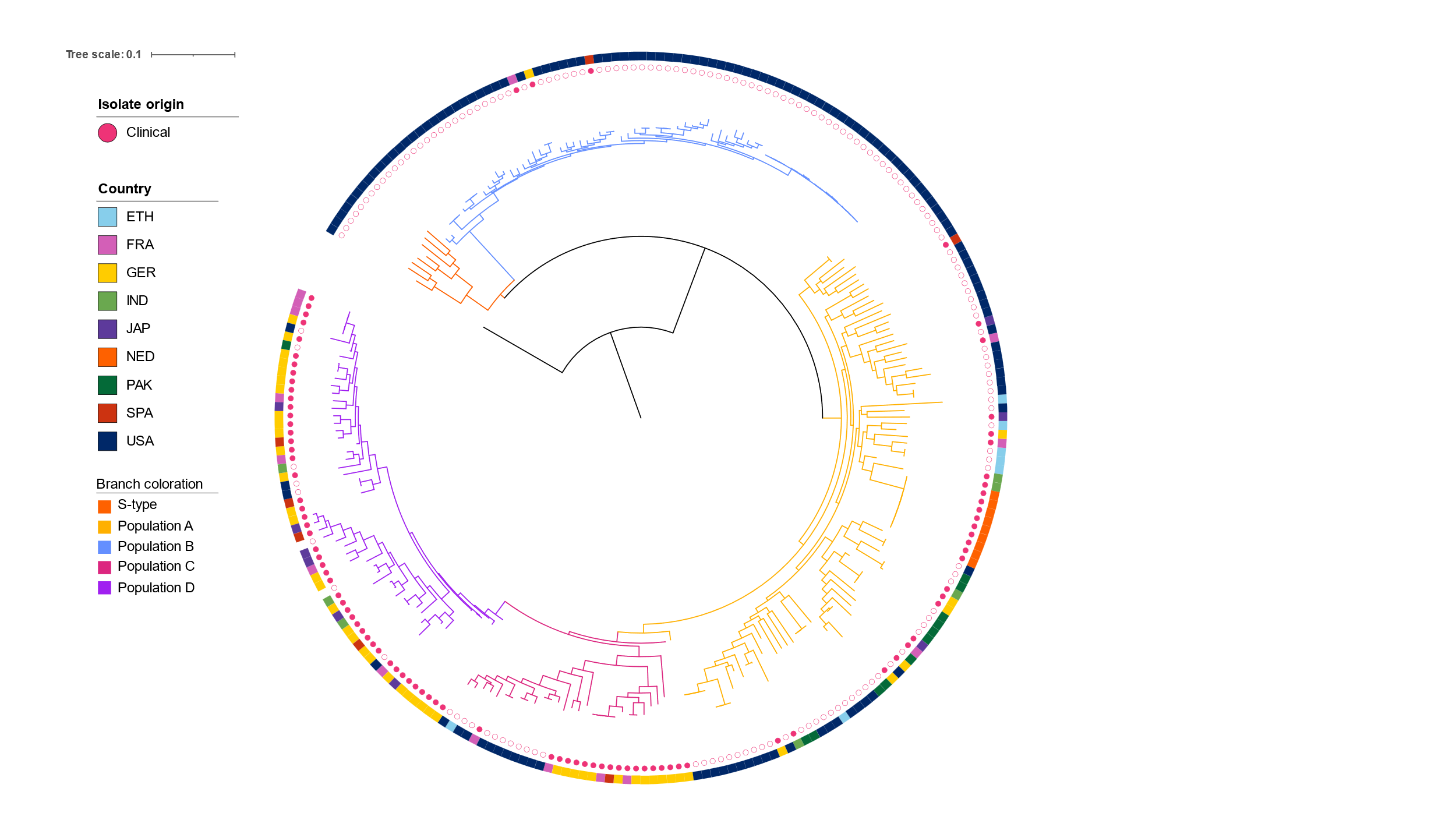

Figure S6. Genome size of *A. flavus* isolates by population as assigned by ancestry coefficient and discriminant analysis of principal components. Asterisks indicate population means were statistically significant (ordinary one-way ANOVA with multiple comparisons; p < 0.05). All other comparisons were nonsignificant. Outliers outside the 5-95 percentile range are indicated as black dots.

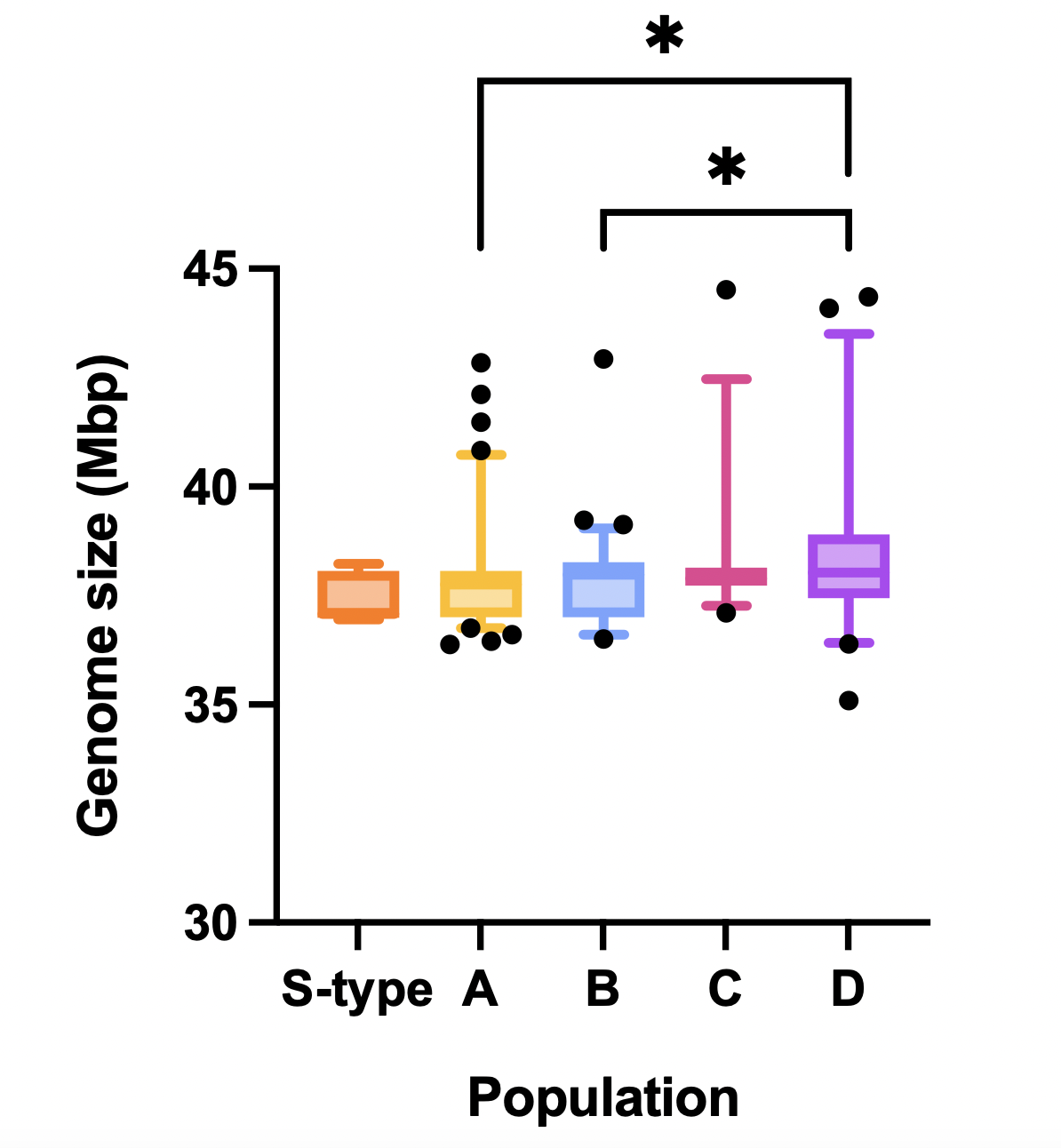

Figure S7. Principal coordinates analysis (PCoA) based on presence or absence of 10,161 orthogroups from the accessory genome. Each circle represents an isolate of *A. flavus*. Isolates are color-coded based on population assignment according to clustering in the discriminant analysis of principal components.

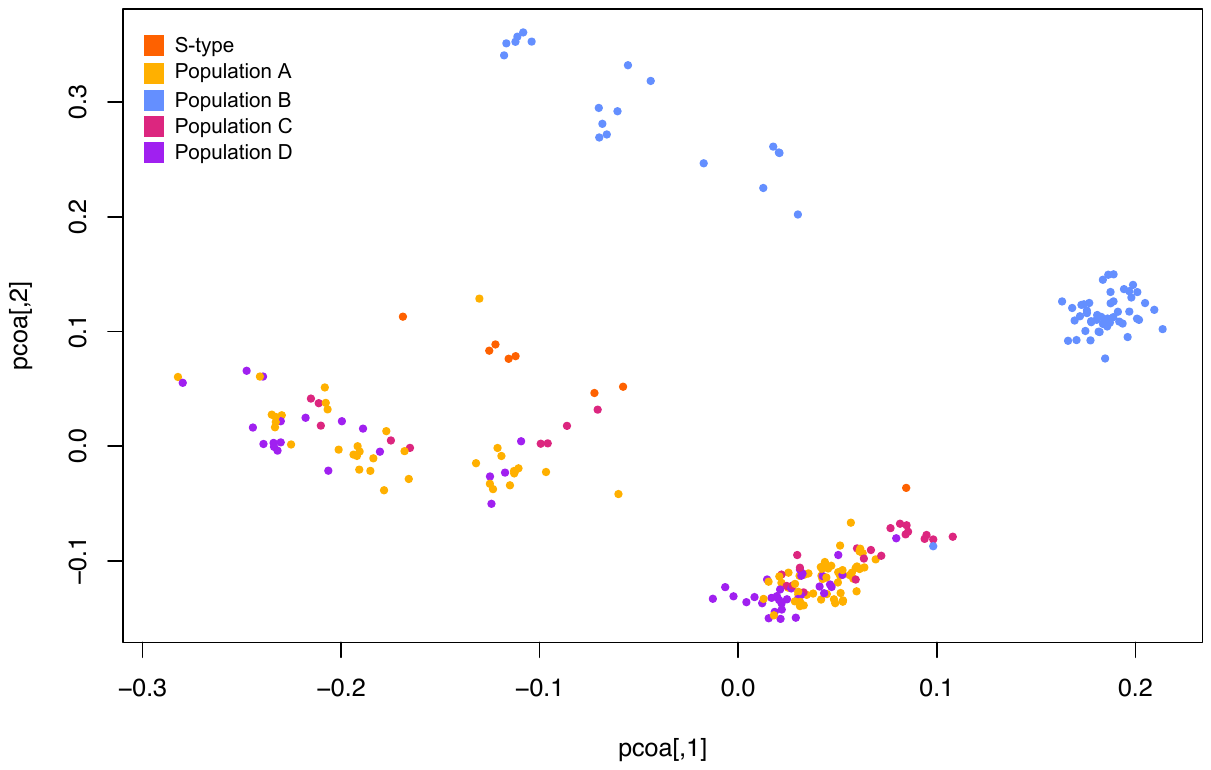

Figure S8. Boxplots of gene ontology (GO) terms more prevalent in population D than other populations of *Aspergillus flavus*. The Y axis represents the number of genes containing the GO term.

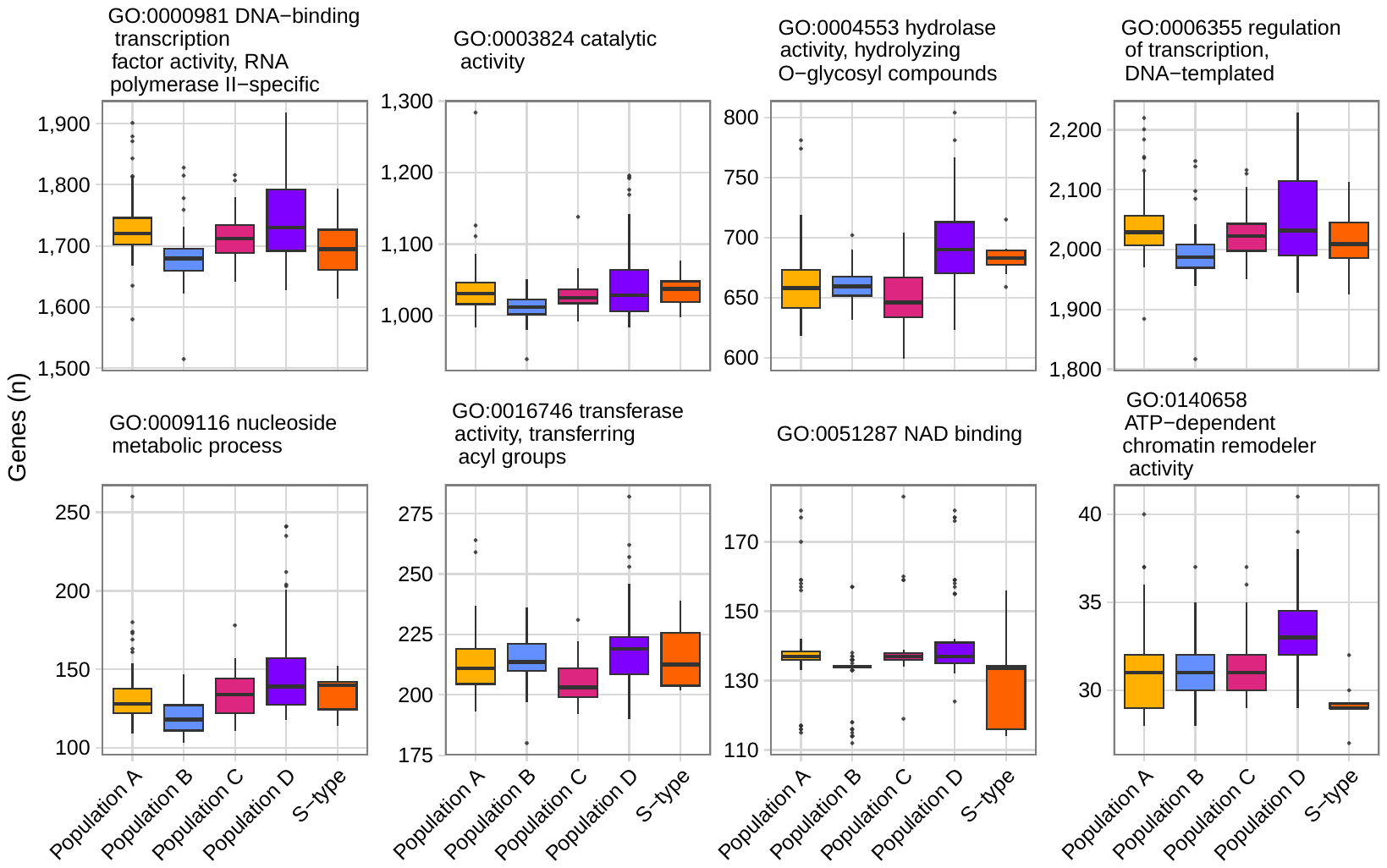

Figure S9. Aflatoxin biosynthetic gene cluster and backbone gene presence and metabolite production across the phylogeny of *A. flavus*. Aflatoxin production data was taken from previously published reports. Clinical isolates are represented by filled in pink circles in the innermost track.

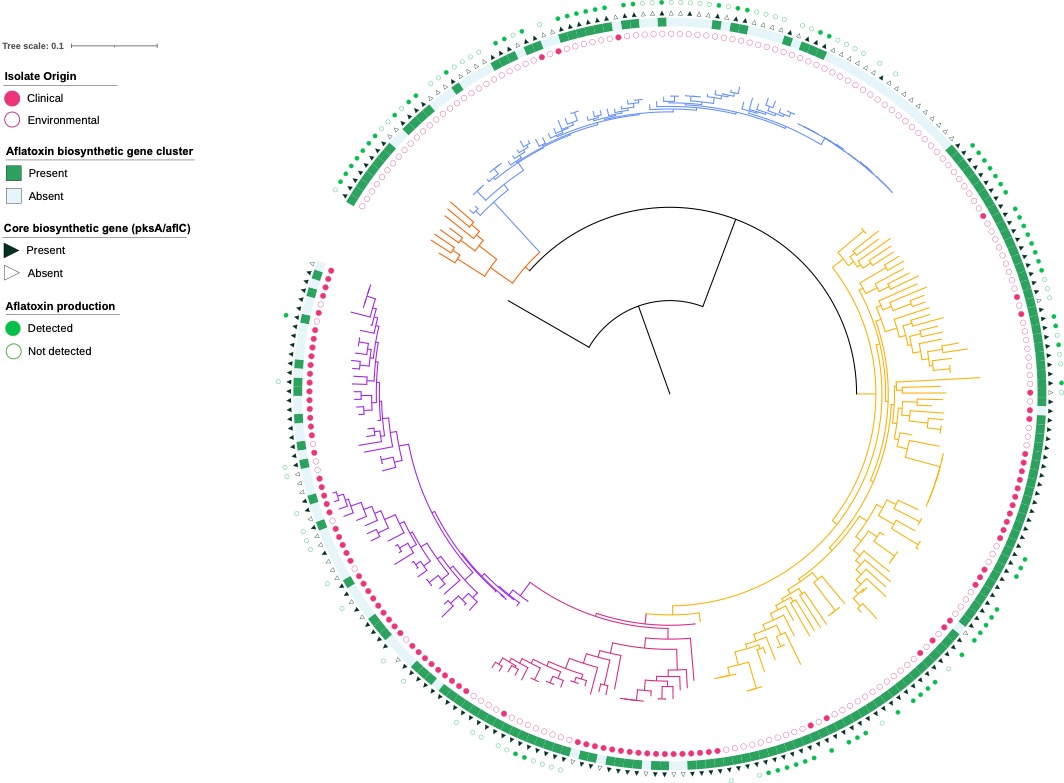

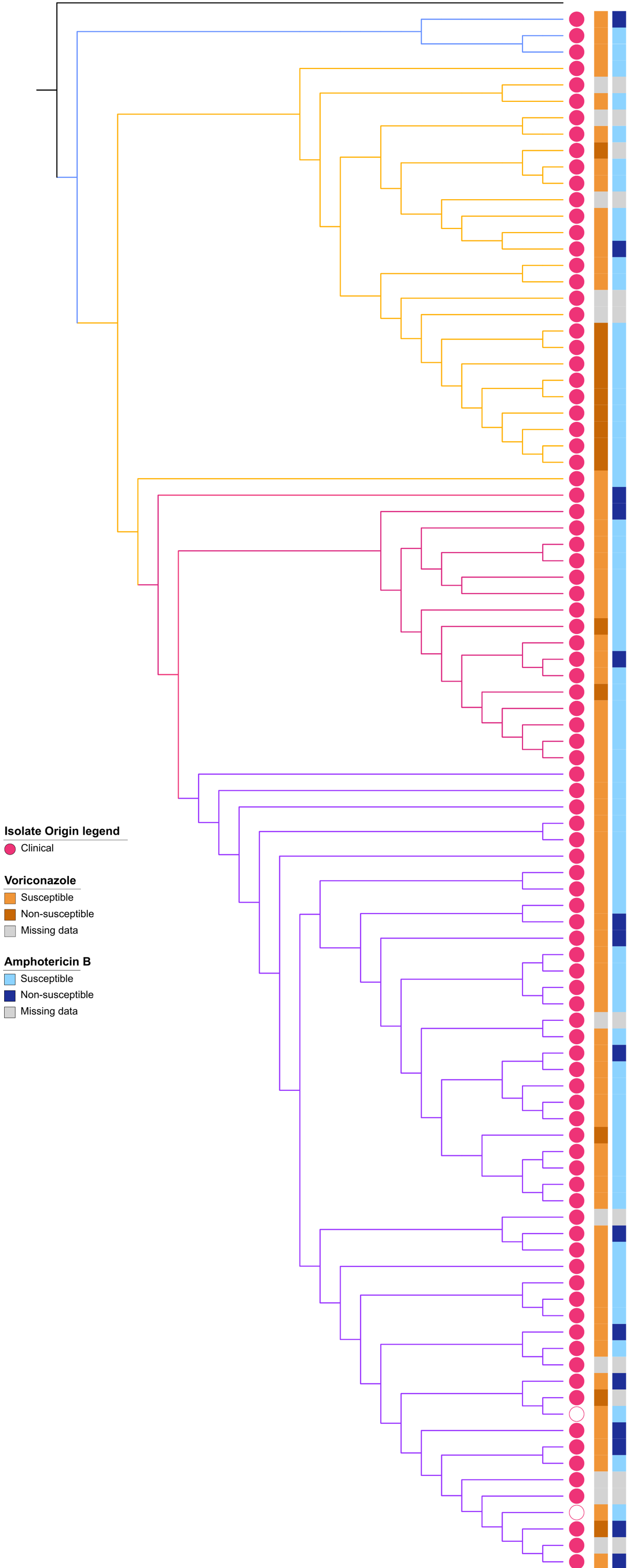
Figure S10. Cladogram with antifungal susceptibility for voriconazole (orange) and amphotericin B (blue) for all isolates with available data for the minimum inhibitory concentration for either antifungal compound. Darker colors indicate MICs above the cutoff (non-susceptible). Lighter colors indicate MICs below the cutoff (susceptible).

Clinical isolates are indicated with filled-in pink circles. Environmental isolates are indicated with empty pink circles. Cladogram branches are color-coded by DAPC population assignment.
